# Reference-based phasing using the Haplotype Reference Consortium panel

**DOI:** 10.1101/052308

**Authors:** Po-Ru Loh, Petr Danecek, Pier Francesco Palamara, Christian Fuchsberger, Yakir A Reshef, Hilary K Finucane, Sebastian Schoenherr, Lukas Forer, Shane McCarthy, Goncalo R Abecasis, Richard Durbin, Alkes L Price

## Abstract

Haplotype phasing is a fundamental problem in medical and population genetics. Phasing is generally performed via statistical phasing within a genotyped cohort, an approach that can attain high accuracy in very large cohorts but attains lower accuracy in smaller cohorts. Here, we instead explore the paradigm of reference-based phasing. We introduce a new phasing algorithm, Eagle2, that attains high accuracy across a broad range of cohort sizes by efficiently leveraging information from large external reference panels (such as the Haplotype Reference Consortium, HRC) using a new data structure based on the positional BurrowsWheeler transform. We demonstrate that Eagle2 attains a ≈20x speedup and ≈10% increase in accuracy compared to reference-based phasing using SHAPEIT2. On European-ancestry samples, Eagle2 with the HRC panel achieves >2x the accuracy of 1000 Genomes-based phasing. Eagle2 is open source and freely available for HRC-based phasing via the Sanger Imputation Service and the Michigan Imputation Server.

---

Haplotype phasing is a central problem in human genetics [1]. Over the past decade, phasing has most commonly been performed via statistical methods applied within a genotyped cohort [2–14]. Wet-lab technologies for direct phasing have also generated considerable recent interest, but these methods are currently much less scalable [15]. In general, the accuracy of statistical phasing methods increases steadily with sample size due to improved modeling of linkage disequi-lbrium and increasing prevalence of identity-by-descent. We and others have recently developed methods that achieve very high statistical phasing accuracy in cohorts comprising a large fraction of a population [8] or containing >100,000 samples [13,14]. However, for smaller cohorts, accuracy of cohort-based statistical phasing is fundamentally limited by the quantity of data available.

Here, we explore an alternative paradigm, reference-based phasing, which can achieve high accuracy even in smaller cohorts by leveraging information from an external reference panel. This paradigm targets a user group complementary to recent methods for phasing very large cohorts [13, 14]. In particular, methods for mapping molecular QTLs using allele-specific reads require accurate phasing information, but recent papers introducing these methods have reported that inaccurate phasing currently limits their potential [16,17].

We present a new reference-based phasing algorithm, Eagle2, which we have incorporated into the Sanger Imputation Service and the Michigan Imputation Server to perform free reference-phasing using the 32,470-sample Haplotype Reference Consortium (HRC) [18]. This approach achieves >2x; improved phasing accuracy over 1000 Genomes-based phasing on small European-ancestry cohorts, with smaller improvements for larger cohort sizes. The Eagle2 algorithm represents a substantial computational advance over existing reference-based phasing algorithms: Eagle2 achieves a 20* speedup over SHAPEIT2 [12]—i.e., genome-wide phasing in 1.5 minutes per sample—with a 10% improvement in accuracy across a range of ancestries. Eagle2 achieves this performance via two key ideas that distinguish it from previous phasing algorithms [3–14]: a new data structure based on the positional Burrows-Wheeler transform [19] and a rapid search algorithm that explores only the most relevant phase paths through a hidden Markov model (HMM). We have released Eagle2 as open-source software (see URLs).

## Results

### Overview of methods

The Eagle2 phasing algorithm takes as input a diploid target sample and a library of reference haplotypes. The statistical model underlying Eagle2 is a haplotype copying model similar to the Li-Stephens model [20] used by previous HMM-based methods. However, Eagle2 has two key differences compared to previous HMM-based methods. First, whereas previous approaches approximate the haplotype structure (e.g., by merging haplotypes into local clusters) to produce a more tractable HMM, Eagle2 efficiently represents the full haplotype structure in a way that losslessly condenses locally matching haplotypes. Second, using this representation, Eagle2 selectively explores the space of diplotypes—i.e., complementary pairs of phased haplotypes—in a way that only expends computation on the most likely phase paths (i.e., diplotypes with highest posterior probabilities). This approach is distinct from the dynamic programming or sampling methods employed by previous phasing software and enables much greater computational efficiency. In more detail, Eagle2 efficiently represents haplotype structure by introducing a new data structure, the HapHedge, which can be generated in linear time using the positional Burrows-Wheeler transform (PBWT) [19]. Eagle2 then explores diplotypes using a branching-and-pruning beam search. We provide a schematic of the method in Figure 1 and present full details in Online Methods and the Supplementary Note.

**Figure 1.**
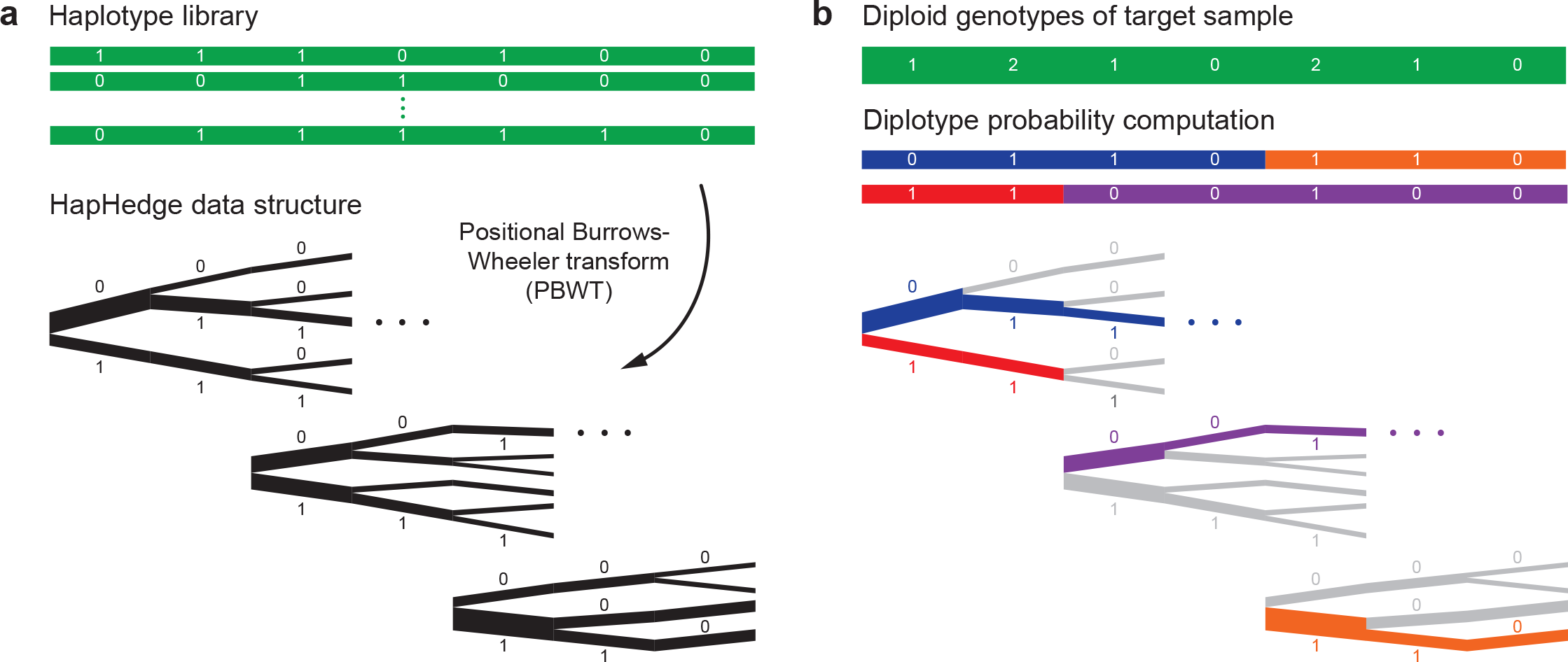
Schematic of the Eagle2 core phasing algorithm. Given diploid genotypes from a target sample along with a haploid reference set of conditioning haplotypes, our algorithm proceeds in two steps. (a) We use the positional Burrows-Wheeler transform [19] to generate a “hedge” of haplotype prefix trees rooted at markers spaced across the chromosome. These trees encode haplotype prefix frequencies, represented here with branch thicknesses. (b) We explore a small set of high-probability diplotypes (i.e., complementary pairs of phased haplotypes), estimating diplotype probabilities under a haplotype copying model by summing over possible recombination points. For each possible choice of recombination points, the HapHedge data structure allows rapid lookup of haplotype segment frequencies. (This illustration is meant to provide intuition for the overall approach; our optimized software implementation first “condenses” reference haplotypes based on the target genotypes. Details are provided in Supplementary Fig. 1 and the Supplementary Note.)

We note that the Eagle2 algorithm is very different from the long-range phasing algorithm we recently developed for phasing extremely large cohorts [13]. (We refer to the previous method as Eagle1.) The basic idea of Eagle1 was to harness identity-by-descent among distant relatives— which is pervasive at very large sample sizes but rare among smaller numbers of samples—to rapidly call phase using a fast scoring approach. In contrast, Eagle2 analyzes a full probabilistic model similar to the diploid Li-Stephens model used by previous HMM-based methods. Consequently, whereas Eagle1 suffered decreased accuracy compared to HMM-based methods when used to phase <50,000 samples, Eagle2 achieves improved accuracy over previous methods for both small and large haplotype reference panel sizes, as we demonstrate below. We note that when a reference panel contains fewer than twice as many samples as the target cohort, Eagle2 iteratively augments the reference panel with inferred target haplotypes (Online Methods); under this paradigm, reference-based phasing should always improve accuracy over cohort-based phasing. We also note that the Eagle1 algorithm was originally only implemented for cohort-based phasing; in this work, we have extended the implementation to reference-based phasing for the sake of comparison. Likewise, we have implemented a cohort-based version of Eagle2 that we also benchmark below.

### Phasing performance using genotyped reference panels

We first benchmarked Eagle2 against previous reference-based phasing methods using reference panels generated by phasing subsets of genotyped cohorts. These benchmarks allowed us to explore a greater range of reference panel sizes and genetic ancestries than currently available in sequenced reference panels (as the *N*=32,470 samples currently in the HRC are predominantly European), understanding that genotyped reference panels containing a limited set of markers are not broadly useful for reference-based phasing. We performed benchmarks using a total of five data sets: the UK Biobank cohort [21] and the four GERA sub-cohorts, which were genotyped on four distinct European, African, East Asian, and Latino genotyping arrays [22,23]. All five data sets were typed on arrays containing 650K–850K autosomal markers with typical heterozygosity and missingness rates, and each data set contained a small subset of mother-father-child trios (Online Methods and Supplementary Table 1).

For the UK Biobank reference-based phasing benchmarks, we generated simulated reference panels by randomly selecting *N*_ref_= 15,000, 30,000, 50,000, or 100,000 samples (not containing trio members) and phasing them using Eagle1 [13]. We phased each subset independently (rather than phasing all samples together and then extracting subsets) to better reflect the phase inaccuracy that would be present in a real reference panel of a given size. We then benchmarked the computational cost and accuracy of reference-based phasing methods by using each panel of 2*N*_ref_ haplotypes to phase sets of other UK Biobank target samples including the 70 European-ancestry trio children, which we used for benchmarking accuracy (Online Methods). To cover a wide range of linkage disequilibrium structure, we performed these benchmarks on chromosomes 1, 5, 10, 15, and 20 (a total of 174,595 markers comprising ≈25% of the genome) using Eagle2, SHAPEIT2 [12], SHAPEIT2 with its `--no-mcmc` option (which increases speed at the expense of accuracy), and a reference-based version of Eagle1 that we implemented for comparison. We also attempted to benchmark Beagle v4.1 [24] but found it was too slow for this benchmark to be practical: for the smallest analysis (chromosome 20 with *N*_ref_=15,000 and *N*_target_=72), Beagle v4.1 required 3.6 days (in contrast to 1.1 minutes for Eagle2). (We note that the focus of Beagle v4.1 [24] is its haploid imputation algorithm, which is much faster than its phasing algorithm.) We did not benchmark HAPI-UR [11] as HAPI-UR does not implement reference-based phasing.

We observed that Eagle2 achieved 12–38x speedups over SHAPEIT2 for performing reference-based phasing using panels of size *N*_ref_=15,000–100,000 (fig. 2a and Supplementary Tables 2 and 3). Moreover, unlike the other methods we benchmarked, the computation time Eagle2 required to phase each target sample was nearly independent of the reference size. (For very large reference panels with *N*_ref_ ≫ 100,000, the computational cost of Eagle2 will eventually increase with *N*_ref_; see Online Methods and the Supplementary Note.) Eagle2 achieved running times similar to Eagle1 and ≈2x faster than SHAPEIT2 `--no-mcmc` (both of which are much less accurate methods than Eagle2 and SHAPEIT2 when used with reference panels of these sizes; see below). All methods had low memory costs (<7GB for *M*=57,753 SNPs on chromosome 1 with *N*_ref_=100,000; Supplementary Table 3).

**Figure 2.**
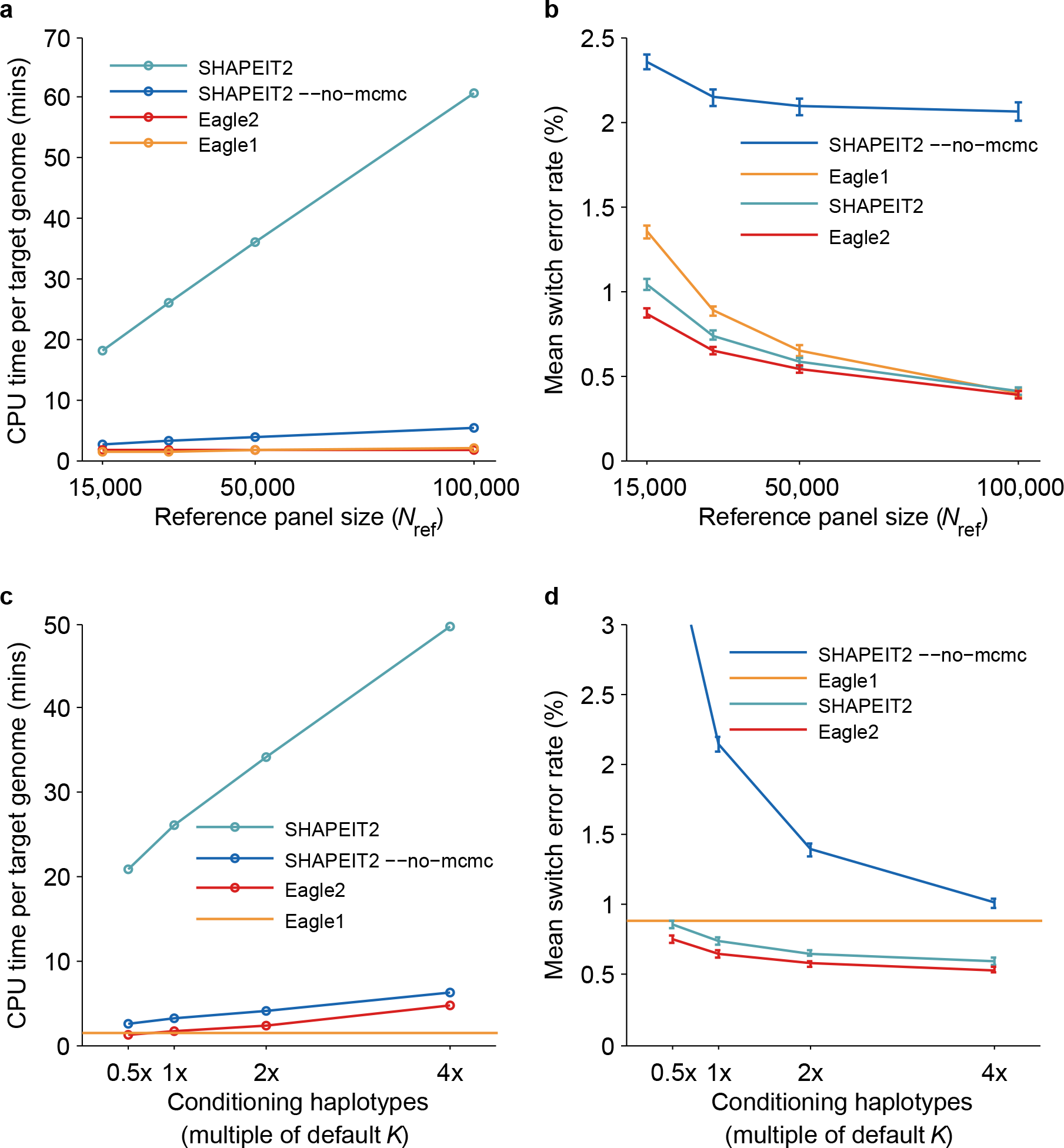
Running time and accuracy of reference-based phasing in UK Biobank benchmarks. We benchmarked Eagle2 and other available methods by phasing UK Biobank trio children using a reference panel generated from Nref = 15,000, 30,000, 50,000, or 100,000 other UK Biobank samples. (a) CPU time per target genome on a 2.27 GHz Intel Xeon L5640 processor.(We analyzed a total of 174,595 markers on chromosomes 1, 5, 10, 15, and 20, representing ≈25% of the genome, and scaled up running times by a factor of 4; see Supplementary Table 3 for details.) (b) Mean switch error rate over 70 European-ancestry trios; error bars,s.e.m. (c, d) CPU time and mean switch error rate as a function of the number of conditioning haplotypes used by SHAPEIT2 and Eagle2 (relative to the default values of *K*=100 and 10,000, respectively). Eagle1 does not have such a parameter, so we display its performance as a horizontal line. Numeric data and additional benchmarks varying the number of conditioning haplotypes used with *N*ref = 15,000, 50,000, and 100,000 are provided in Supplementary Table 2.

In our accuracy benchmarks, which we computed using gold standard trio phase calls, we observed that Eagle2 achieved 5–16% lower switch error rates [2] compared to SHAPEIT2, with larger gains for lower values of *N*_ref_ in the 15,000–100,000 range (Fig. 2b and Supplementary Table 2). Eagle2 achieved 4–36% lower switch error rates than Eagle1 and 63–81% lower switch error rates than SHAPEIT2 `--no-mcmc` in the same *N*_ref_ range. All differences were statistically significant (binomial *p*<0.003 for each comparison of Eagle2 with SHAPEIT2, *p*<0.03 for each comparison with Eagle1, and *p*<10^−21^ for each comparison with SHAPEIT2 `--no-mcmc`).

Both Eagle2 and SHAPEIT2 have an important parameter, *K*, that specifies the number of conditioning haplotypes used to phase each target sample and thus adjusts the speed-accuracy tradeoff. We therefore also investigated the effects of varying *K*. (We note that the default values and precise meaning of this parameter are different for Eagle2 vs. SHAPEIT2; by default, SHAPEIT2 locally selects *K*=100 best reference haplotypes in each 2Mb window, while Eagle2 selects a fixed set of *K*=10,000 best reference haplotypes to use for the entire chromosome. This difference may be responsible for the slightly lower rate of improvement in accuracy of Eagle2 relative to that of SHAPEIT2 as *N*_ref_ increases at fixed *K* in Fig. 2b.) We considered a range of values of K from 0. 5–4 times the default *K*, similar to previous benchmarks of SHAPEIT2 [12]. The effects of varying *K* were broadly consistent for Eagle2, SHAPEIT2, and SHAPEIT2 –-no-mcmc: all methods required similarly increased computation time and achieved improved accuracy with larger values of *K* (Fig. 2c,d and Supplementary Tables 2 and 3). In particular, increasing the number of conditioning haplotypes by a factor of 4* required 2–3* more computation time for both Eagle2 and SHAPEIT2 while achieving similar decreases in switch error rates (12–20% and 17–19%, respectively, for *N*_ref_ =15,000–100,000; Supplementary Table 2). All improvements were statistically significant (binomial *p*<10^−7^).

To assess the robustness of these accuracy benchmarks across genetic ancestries, we performed a similar set of benchmarks using the European, African, East Asian, and Latino GERA subcohorts (Online Methods). Because the latter three sub-cohorts were relatively small (Supplementary Table 1), we generated a single simulated reference panel for each sub-cohort containing all samples not belonging to trio pedigrees (*N*_ref_=3,817, 5,164,7,144, and 61,684 for the African, East Asian, Latino, and European sub-cohorts). We phased the three smaller panels using SHAPEIT2 and phased the European panel using Eagle1. We then benchmarked reference-based phasing accuracy by phasing the trio parents within each sub-cohort using the panel generated from that sub-cohort, running each method with default parameter settings. (We phased trio parents rather than trio children for these benchmarks because the three smaller data sets contained only 3–7 independent trios each; Supplementary Table 1.) These benchmarks confirmed our findings from the UK Biobank data: Eagle2 achieved 5–23% lower switch error rates than SHAPEIT2, and we observed the same relative ordering of accuracies as before across all sub-cohorts (Figure 3 and Supplementary Table 4). All differences were statistically significant (binomial *p*<10^−7^). We note that every method had a higher switch error rate in the GERA European sub-cohort compared to the UK Biobank, presumably due primarily to a more diverse set of ancestries represented. In general, absolute switch error rates are not directly comparable among data sets due to differences in demography and genotyping properties (e.g., chip density, allele frequency distribution, and genotype error rate).

**Figure 3.**
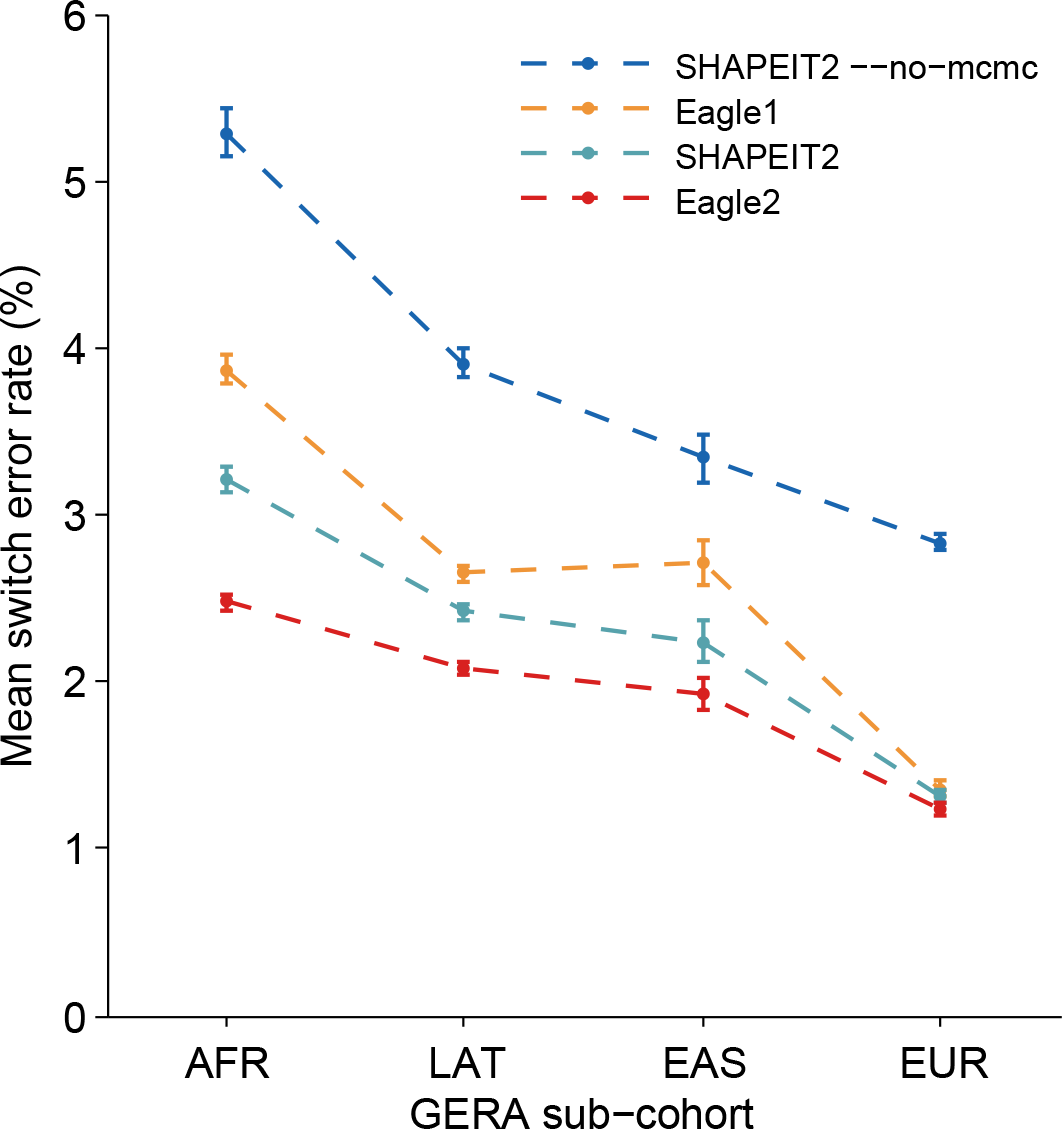
Accuracy of reference-based phasing in GERA benchmarks. We phased trio parents in each GERA sub-cohort using a reference panel generated from all other non-familial samples in the same sub-cohort. We ran each method with default parameter settings on all 22 autosomes and computed aggregate mean switch error rates; error bars, s.e.m. Standard errors for the European-ancestry sub-cohort are over 400 parent samples. Standard errors for the other three sub-cohorts are over 25 SNP blocks. Numeric data and additional benchmarks varying the number of conditioning haplotypes used by each method are provided in Supplementary Table 4.

### Phasing accuracy using the 1000 Genomes and HRC reference panels

We next benchmarked reference-based phasing using either the 1000 Genomes Project Phase 3 reference panel (containing *N*=2,504 samples from 26 populations) [25] or the HRC reference panel r1.1 (containing *N*=32,470 samples mostly of European ancestry) [18]. For these benchmarks, we used 1000 Genomes trio children as target samples, removing all 1000 Genomes trios from each reference panel before running the analyses. We phased chromosome 1, and to emulate typical genotyping density, we restricted the SNP set to 31,853 sites typed on 23andMe (customized Illumina) chips.

Given the predominantly European composition of the HRC panel, the benchmarks on 32 CEU trio children were of primary interest, and we observed in these benchmarks that all methods achieved substantially improved accuracy using the HRC panel versus the 1000 Genomes panel (Figure 4 and Supplementary Table 5). For each choice of reference panel, Eagle2 achieved the lowest switch error rate, consistent with our previous results. For phasing small European cohorts, Eagle2 with the HRC panel provides a >2x improvement in accuracy over 1000 Genomes-based phasing: 1.35% (s.e. 0.04%) switch error rate versus 3.52% (0.06%) using SHAPEIT2 or 3.27% (0.06%) using Eagle2 with the 1000 Genomes reference (Figure 4 and Supplementary Table 5).

**Figure 4.**
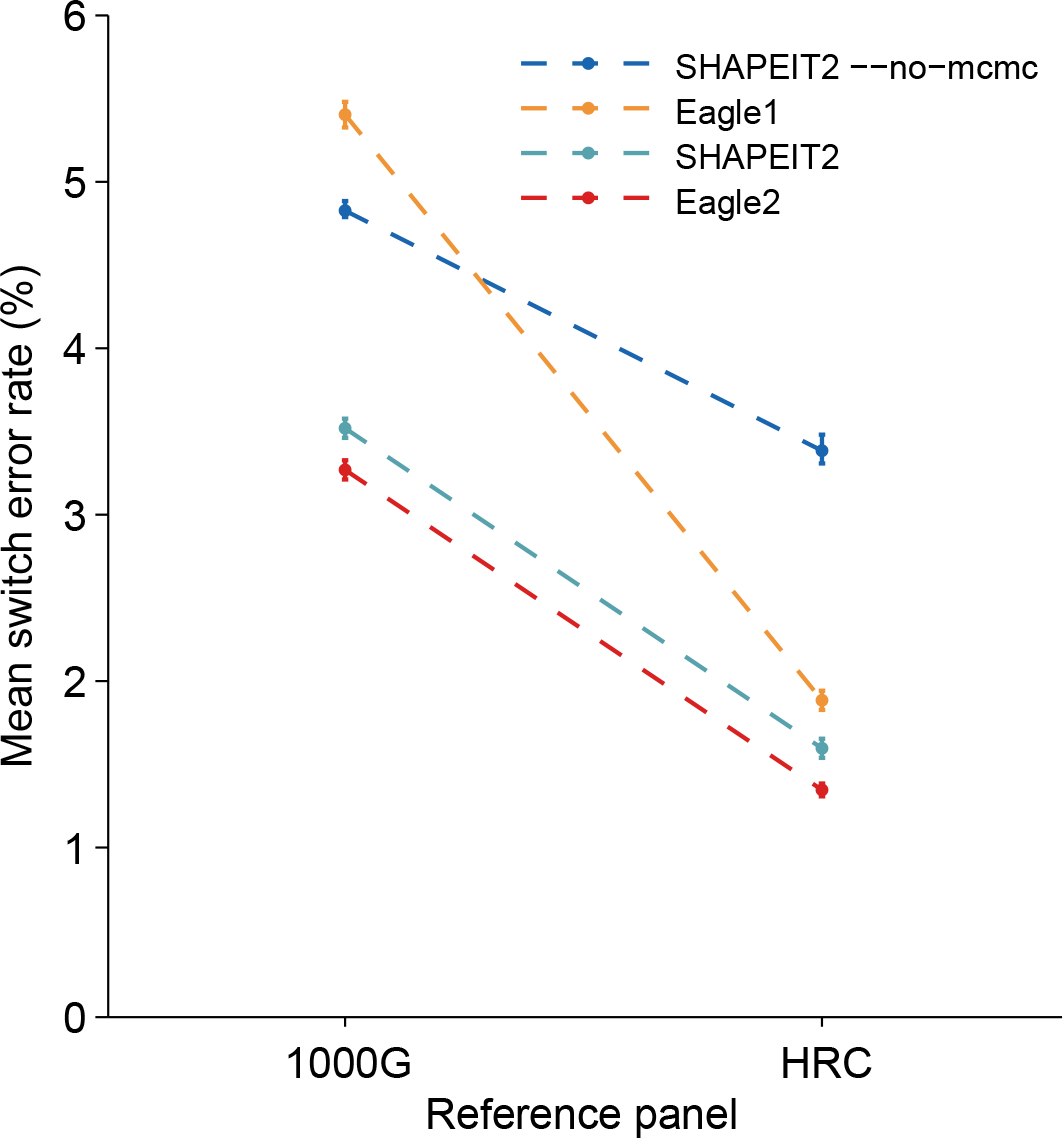
Accuracy of reference-based phasing using the 1000 Genomes and HRC panels. We phased 32 trio children from the 1000 Genomes CEU population using either the 1000 Genomes Phase 3 reference panel or the Haplotype Reference Consortium panel (excluding trios in either case). We analyzed chromosome 1, and to emulate a typical use case, we restricted the data to 31,853 markers (genotyped on 23andMe chips). We plot mean switch error rates; error bars, s.e.m. over samples. Numeric data and additional benchmarks on other 1000 Genomes populations are provided in Supplementary Table 5.

We also benchmarked accuracy in all other 1000 Genomes populations containing >1 trio. We phased trio children in 31 Han Chinese (CHS) trios, 30 Peruvian (PEL) trios, 15 Punjabi (PJL) trios, and 19 Yoruba (YRI) trios using either the 1000 Genomes panel or the HRC panel, and we observed that in all cases Eagle2’s accuracy was either slightly better or statistically indistiguish-able from SHAPEIT2's (Supplementary Table 5). Specifically, the differences between Eagle2 and SHAPEIT2 were not significant for PEL with either reference panel and for YRI with HRC; all other differences were significant (binomial *p*<0.04). Interestingly, all methods achieved lower accuracy using the HRC panel versus the 1000 Genomes panel (Supplementary Table 5). Given that the HRC panel contains the 1000 Genomes panel, this observation suggests that the inclusion of ≈30,000 additional predominantly European samples reduced the ability of each method to model the haplotype structure of non-European populations. However, we did not observe this phenomenon when phasing the two non-European UK Biobank trios using increasing numbers of European reference haplotypes (Supplementary Table 6), so this observation may be specific to the current HRC release (r1.1); development of the HRC is ongoing.

### Phasing performance without a reference panel

Lastly, we assessed the performance of Eagle2 when applied to cohort-based phasing, which we also implemented in our software. The Eagle2 cohort-based phasing algorithm starts by running the first two steps of Eagle1 [13] to rapidly produce rough haplotype estimates and then refines these estimates using the Eagle2 core phasing algorithm (Online Methods). We benchmarked Eagle2, Eagle1, and SHAPEIT2 on subsets of the UK Biobank data set containing *N* = 5,000, 15,000, 50,000, or 150,000 samples (including trio children and excluding trio parents). We phased chromosomes 1, 5, 10, 15, and 20 as in our UK Biobank reference-based phasing benchmarks, and we allowed each computational job up to 5 days to complete. We observed that Eagle2 exhibited computational efficiency similar to Eagle1, achieving 5-6x speedups over SHAPEIT2 in the analyses SHAPEIT2 was able to complete (*N*=5,000 and *N*=15,000) (Fig. 5a and Supplementary Tables 7 and 8). Eagle2 also exhibited close-to-linear run time scaling across this sample size range, breaking even with Eagle1 at *N*≈30,000 and achieving faster running times for larger *N*.

**Figure 5.**
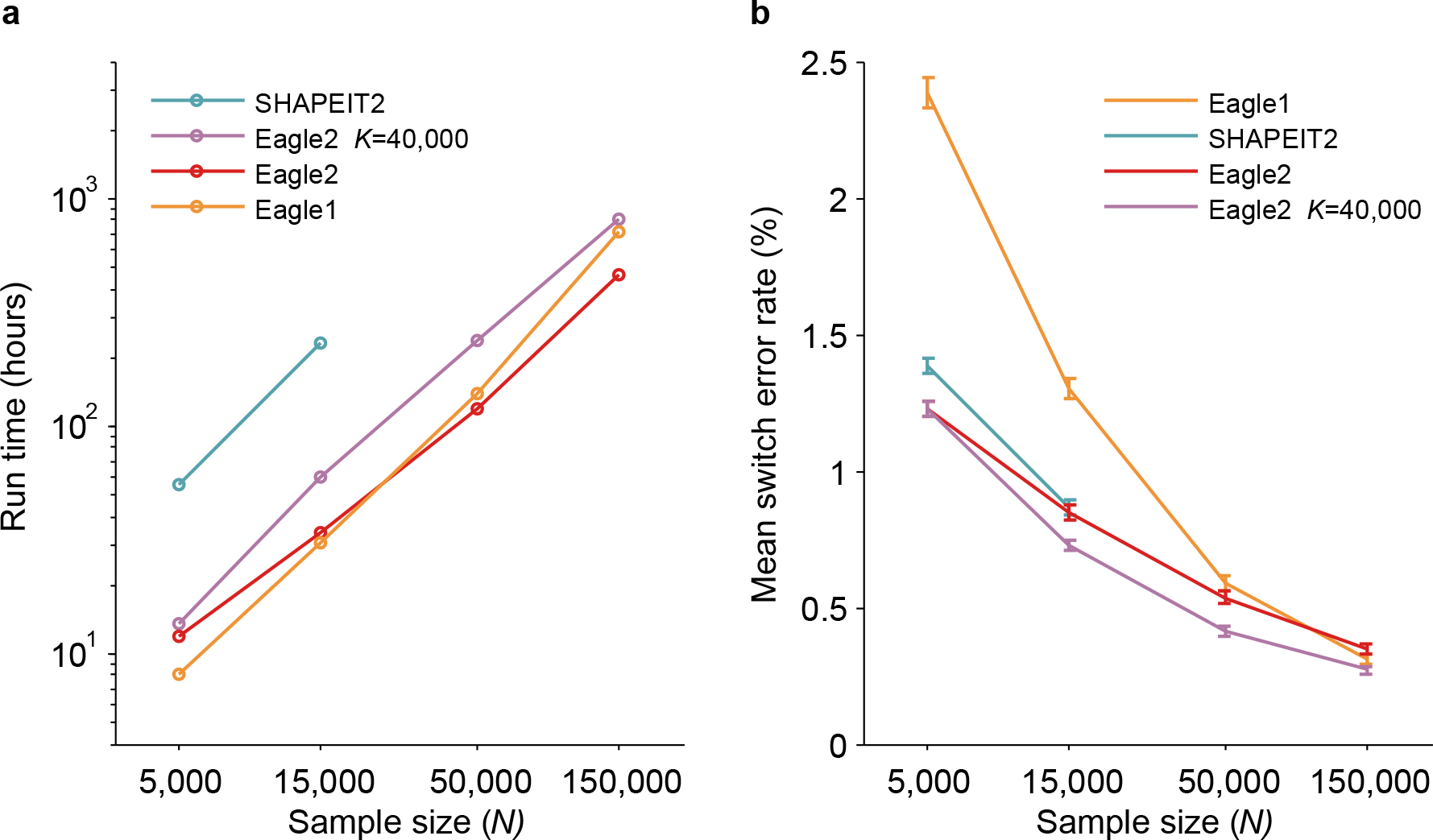
Running time and accuracy of cohort-based phasing in the UK Biobank cohort. We benchmarked Eagle2 and other available phasing methods on N=5,000, 15,000 50,000, and150,000 UK Biobank samples (including trio children and excluding trio parents). (a) Total wall clock time for genome-wide phasing on a 16-core 2.60 GHz Intel Xeon E5-2650 v2 processor. (We analyzed a total of 174,595 markers on chromosomes 1, 5, 10, 15, and 20, representing ≈25% of the genome, and scaled up running times by a factor of 4; see Supplementary Table 8 for per-chromosome data.) SHAPEIT2 was unable to complete the *N*=50,000 chr1 and chr5 analyses and was uanble to complete any of the *N*=150,000 analyses in 5 days, the run time limit for single compute jobs. (b) Mean switch error rate over 70 European-ancestry trios; error bars,s.e.m. Numeric data and additional benchmarks varying the number of conditioning haplotypes used by Eagle2 are provided in Supplementary Table 7.

In our accuracy benchmarks using the 70 European-ancestry UK Biobank trios, we observed that Eagle2 achieved better accuracy than SHAPEIT2 and Eagle1 for *N*≤50,000, as expected (Fig. 5b and Supplementary Tables 7 and 8). At *N*=150,000, Eagle1 achieved a slightly lower switch error rate (0.31%, s.e. 0.02%) than Eagle2 (0.35%, s.e. 0.02%). However, we observed that running Eagle2 with 4x the default number of conditioning haplotypes (i.e., K=40,000) achieved the lowest error rates across all sample sizes tested (0.27%, s.e. 0.02% at *N*=150,000). Both differences were statistically significant (binomial *p*<0.0006). Finally, we confirmed that Eagle2 achieved better phasing accuracy than SHAPEIT2 or Eagle1 when used to phase the GERA samples within each GERA sub-cohort (Supplementary Table 9), with switch error rates consistent with our earlier reference-based benchmarks (Figure 3 and Supplementary Table 4). All differences were statistically significant (binomial *p*<0.002).

## Discussion

We have described a new phasing algorithm, Eagle2, which we have incorporated into the Sanger Imputation Service and the Michigan Imputation Server to offer free reference-based phasing using the *N*=32,470-sample Haplotype Reference Consortium panel. This service enables high-accuracy phasing even in smaller cohorts, which was not previously possible. Eagle2 achieves substantial gains in speed and accuracy over previous methods via a novel search-based algorithm employing the positional Burrows-Wheeler transform. We believe this method is timely, as large sequenced reference panels (e.g., the HRC) are now becoming available for use— but must be utilized via analyses run on central servers due to consent restrictions. We anticipate that Eagle2's phasing speed—1.5 minutes per genotyped sample— will help keep computation tractable as demand for this service increases. Additionally, we anticipate that our release of Eagle2 as open-source software will aid in future method development and integration into analysis pipelines.

We note that Eagle2 targets a distinct user group compared to very recent work on phasing very large cohorts [13,14]. In particular, our Eagle1 method [13] is targeted at phasing very large (*N*> 100,000) cohorts and achieves much lower accuracy than both Eagle2 and previous methods when used to phase smaller cohorts. The SHAPEIT3 method [14] is likewise targeted at phasing “biobank scale datasets.” We were unable to benchmark SHAPEIT3 given that the SHAPEIT3 software has not yet been released, but ref. [14] indicates that its primary advance is removing a quadratic complexity component of the SHAPEIT2 algorithm that becomes significant as *N* increases beyond 10,000 samples; this computational speedup comes at the expense of reduced accuracy. The benchmarks in ref. [14] suggest that if used to perform HRC-based phasing at N_ref_=32,470, SHAPEIT3 would be ≈3x faster but roughly 20% less accurate than SHAPEIT2; in contrast, Eagle2 is ≈20x faster and ≈10% more accurate than SHAPEIT2 at this sample size. (In practice, the SHAPEIT license precludes its use for reference-based phasing on the Sanger and Michigan HRC servers.)

While we believe that reference-based phasing using large reference panels such as the HRC is a valuable phasing paradigm, we note a few limitations. First, reference-based phasing accuracy is limited not only by reference panel size but also by genotyping and phasing accuracy in the reference panel. In particular, the HRC reference haplotypes are largely derived from low-coverage sequencing data (which is generally prone to higher errors in genotype calling), and efforts to improve the accuracy of the reference panel are ongoing. Second, for reference-based phasing to be effective, the reference panel needs to contain a sizable subset of samples with genetic ancestry well-matched to the target samples. Consequently, phasing using the HRC is currently only advantageous for European-ancestry target samples, although plans are underway to grow the HRC to better represent worldwide populations. Third, for very large cohorts (substantially larger than the reference size), we expect that reference-based phasing will achieve only marginal gains in accuracy over cohort-based phasing. For such cohorts, we expect that cohort-based phasing may remain the preferred option due to ease of execution—although reference sizes are growing, and if the end goal is HRC-based imputation, then we expect that HRC-based pre-phasing [26] will be more convenient. Finally, we have not tuned Eagle2 for phasing sequenced samples [27,28]. While a preliminary benchmark against SHAPEIT2 suggests that Eagle2 achieves gains in speed and accuracy on sequence data comparable to its improvements on genotype data (Supplementary Table 10), more work will be needed to optimize the method and benchmark it against other approaches (e.g., the recent SHAPEITR method [29], which is available via the Oxford Statistics Phasing Server; see URLs). Despite these limitations, we expect that reference-based phasing using Eagle2 and the HRC panel will be a valuable resource providing free, fast, and accurate phasing to the scientific community.

### URLs

Eagle2 software and source code,http://www.hsph.harvard.edu/alkes-price/software/.

**SHAPEIT2 software**,http://mathgen.stats.ox.ac.uk/genetics_software/shapeit/shapeit.html.

**Beagle v4.1 software**,http://faculty.washington.edu/browning/beagle/beagle.html.

**PLINK software**,http://www.cog-genomics.org/plink2.

**UK Biobank**,http://www.ukbiobank.ac.uk/.

**UK Biobank Genotyping and QC Documentation**,http://www.ukbiobank.ac.uk/wp-content/uploads/2014/04/UKBiobank_genotyping_QC_documentation-web.pdf.

**GERA** data set,http://www.ncbi.nlm.nih.gov/projects/gap/cgi-bin/study.cgi?study_id=phs000674.v1.p1.

**1000 Genomes data set**,http://www.1000genomes.org/.

**Haplotype Reference Consortium**,http://www.haplotype-reference-consortium.org/.

**Sanger Imputation Service**,https://imputation.sanger.ac.uk/.

**Michigan Imputation Server**,https://imputationserver.sph.umich.edu/.

**Oxford Statistics Phasing Server**,https://phasingserver.stats.ox.ac.uk/.

## ACKNOWLEDGMENTS

**Acknowledgments.** We are grateful to S. Linderman, N. Patterson, L. O’Connor, A. Gusev, and B. van de Geijn for helpful discussions. This research was conducted using the UK Biobank Resource and was supported by US National Institutes of Health grants R01 HG006399 and R01 MH101244 and US National Institutes of Health fellowship F32 HG007805. PD, SM, RD and the Sanger Institute HRC server are supported by Wellcome Trust grant WT098051. HF was supported by the Fannie and John Hertz Foundation. Computational analyses were performed on the Orchestra High Performance Compute Cluster at Harvard Medical School, which is partially supported by grant NCRR 1S10RR028832-01, and on the Lisa Genetic Cluster Computer (http://www.geneticcluster.org) hosted by SURFsara and financially supported by the Netherlands Scientific Organization (NWO 480-05-003 PI: Posthuma) along with a supplement from the Dutch Brain Foundation and the VU University Amsterdam.

## Online Methods

**Eagle2 core algorithm for phasing a single target sample using a set of reference haplotypes**.
Here we outline the key ideas underlying the Eagle2 core algorithm for phasing a single target sample using a set of reference haplotypes, which has three main steps.

*Step 1: Selection of conditioning haplotypes.* Eagle2 first identifies a subset of *K*=10,000 conditioning haplotypes by ranking reference haplotypes according to the number of discrepancies between each reference haplotype and the homozygous genotypes of the target sample. As in our previous work [13], we perform computation on blocks of up to 64 SNPs at once using bitwise arithmetic; thus, the total computational cost of subset selection is linear in *N_ref_* with a very small constant factor (ignoring time to rank the results, which is negligible in practice). The constant factor is small enough that this step constitutes only a small fraction of the total run time for *N*_ref_<100,000. We note that our discrepancy metric does not make use of inferred phase of the target genotypes (which is possible within an iterative phase refinement scheme) and produces a single set of conditioning haplotypes to use for the entire region being phased, in contrast to the sophisticated approach used by SHAPEIT2 [12]. However, Eagle2 is able to condition on 100x more haplotypes than SHAPEIT2, which we suspect makes selection of conditioning haplotypes much less important. The overall complexity of this step is *O(MN*_ref_) in both time and memory.

*Step 2: Generation of HapHedge data structure.* Eagle2 next generates a HapHedge data structure on the selected conditioning haplotypes. The HapHedge encodes a sequence of haplotype prefix trees (i.e., binary trees on haplotype prefixes) rooted at a sequence of starting positions along the chromosome, thus enabling fast lookup of haplotype frequencies (Figure 1). (In practice, we start a new tree roughly once per heterozygous site in the target sample; Supplementary Fig. 1.) The key features of the HapHedge are linear-time construction, linear-memory representation, and constant-time prefix extension, all with small constant factors. The compact in-memory representation of the HapHedge is achieved via radix trees (Supplementary Fig. 2), while linear-time construction is achieved via the positional Burrows-Wheeler transform [19]. In its simplest form, the PBWT iteratively creates sorted lists of haplotype prefixes, moving the prefix start point from right to left. Our algorithm extends this procedure to convert the lists of sorted prefixes into prefix trees; see the Supplementary Note for details. The overall complexity of this step is *O(MK)* in both time and memory.

*Step 3: Exploration of the diplotype space.*Having prepared a HapHedge of conditioning haplotypes, Eagle2 performs phasing using a statistical model similar to the Li-Stephens haplotype copying model [5,20] used by previous HMM-based methods. However, in contrast to previous methods, Eagle2 applies two new ideas to perform fast and accurate phase inference under this model. The first idea is a new way to efficiently compute haplotype probabilities under a copying model. Naively, such computations require exponential time because of the combinatorial explosion of possible recombination points. The standard approach to overcoming this barrier is to observe that within a *K*_HMM_-state HMM, recursion allows computation of *all* marginal probabilities (for all *K*_hmm_ states at each of M positions) in *O(MK*_HMM_^2^) time. With Eagle2, we take a completely different recursive approach that computes the probability of a *single* haplotype in *O(M)* time--independent of the number of reference haplotypes K--after creation of the HapHedge in *O(MK)* time. The HapHedge essentially consolidates all reference haplotypes sharing a common prefix (starting at any given position) into a single atom of data, thus eliminating future computation that scales with K.

Of course, being able to very rapidly compute the probability of a single haplotype is only useful if we can identify a small subset of haplotype probabilities that are worth computing; to this end, Eagle2 employs a second key idea. We perform a beam search from left to right across the chromosome, propagating a small set of likely diplotypes that represent most of the posterior probability mass in the local diplotype space. This approach essentially focuses computational effort on a small subset of the diplotype space (vs. expending computation evenly across the space as in HMM recursion), which is advantageous when most of the space is probabilistically unfavorable but difficult to discard *a priori.* Full mathematical and engineering details are provided in the Supplementary Note. The overall complexity of this step is *O(MHP*) time and *O(MP*+*HP*) memory, where *H* and *P* are “history length” and “beam width” parameters of the beam search described in the Supplementary Note.

**Eagle2 algorithm for reference-based phasing with multiple target samples.** In practice, reference-based phasing is typically performed on a target set containing many samples, allowing the potential to improve phasing accuracy by using inferred target haplotypes to phase each other. By default, Eagle2 performs a variable number of phasing iterations chosen based on the relative size of the target (*N*_target_) and the reference (*N*_ref_). This behavior is intended to allow Eagle2 to automatically benefit from increased statistical power available from larger target sample sizes. Specifically, if *N*_target_ < *N*_ref_/2, Eagle2 performs only one phasing iteration (phasing each target sample using only the reference haplotypes). If *N*_ref_/2≤*N*_target_<2*N*_ref_, Eagle2 performs two iterations, augmenting the reference panel with the inferred target haplotypes during the second iteration. If *N*_target_≥2*N*_ref_, Eagle2 performs three iterations in an analogous manner. Whenever Eagle2 performs more than one iteration, all iterations prior to the final iteration use K/2 conditioning haplotypes to save time, given that the last iteration has the most impact on accuracy. The number of iterations can also be set directly via the --pbwtlters parameter.

**Eagle2 algorithm for cohort-based phasing**. To perform cohort-based phasing (i.e., without a reference), Eagle2 employs an iteration similar to the above approach, but prior to running two iterations of the Eagle2 core phasing algorithm as described above, it first runs the first two steps of the Eagle1 algorithm, which rapidly detect identical-by-descent segments and use them to call phase [13]. Within small data sets, identity-by-descent is less common, but our results indicate that the two subsequent iterations of the Eagle2 core phasing algorithm are able to rapidly refine phase calls given even an inaccurate set of initial phase calls.

**UK Biobank data set.**We analyzed data from the UK Biobank consisting of 152,729 samples typed at ≈800,000 SNPs. Using PLINK [30] (see URLs), we removed 480 individuals marked for exclusion from genomic analyses based on missingness and heterozygosity filters and 1 individual who had withdrawn consent, leaving 152,248 samples (see URLs, Genotyping and QC). We restricted the SNP set to autosomal, biallelic SNPs with missingness ≤10% and we further excluded 65 autosomal SNPs found to have significantly different allele frequencies between the UK BiLEVE array and the UK Biobank array, leaving 707,524 SNPs (57,753 on chr1, 41,538 on chr5, 34,588 on chr10, 22,367 on chr15, and 18,349 on chr20). We identified 72 trios based on IBS0<0.001, sex of parents, and age of trio members (see URLs, Genotyping and QC). Of the 72 trio children, 69 self-reported British ethnicity, one self-reported Indian ethnicity, and one selfreported Caribbean ethnicity. The remaining trio child did not self-report any ethnicity, but her parents self-reported Irish and “Any other white background” as their ethnicities, so we included this trio child in the 70 European-ancestry trio children we used to benchmark phasing accuracy.

**GERA data set.** We analyzed GERA samples (see URLs; dbGaP study accession phs000674.v1.p1) typed on each of the four GERA ancestry-specific chips (European, African, East Asian, and Latino); QC is described in ref. [22]. We directly analyzed all samples and all autosomal SNPs in each of the four sub-cohorts (Supplementary Table 1). We identified independent trios in each sub-cohort according to pedigree information provided with the data release.

**Phasing software versions and parameter settings.** We benchmarked Eagle1 [13], Eagle2 (v2.1), SHAPEIT v2 (r790) [12], and Beagle v4.1 (22Feb16.8ef) [24] using the Oxford genetic map (supplied with SHAPEIT and Eagle). When running Eagle2 in reference-based phasing mode, we turned off imputation of missing genotypes using the `--noImpMissing` flag; otherwise, we ran all methods using their default parameter settings unless explicitly testing non-default settings. Specifically, the non-default parameter settings we tested were the `--no-mcmc` and `--states` (K) option of SHAPEIT2 and the `--Kpbwt` option of Eagle2.

**Reference-based phasing using genotyped reference panels.**For our reference-based phasing benchmarks using genotyped UK Biobank data, we constructed reference panels by randomly selecting *N*_ref_=15,000, 30,000, 50,000, or 100,000 samples (disjoint from the 72 UK Biobank trios) and phasing these samples using Eagle1 (as phasing using SHAPEIT2 would have required several weeks [13]). We then applied each reference-based phasing method to phase chromosomes 1, 5, 10, 15, and 20 of the *N*_target_=72 trio children using each simulated reference panel, and we compared the phased output against trio phase calls to compute switch error rates [2,13]. In our results, we report mean switch error rates and s.e.m. over the 70 European-ancestry trio children (according to self-reported ethnicity; see above).

To benchmark per-sample computational cost of reference-based phasing, we performed an additional set of analyses in which we phased 1,000 randomly selected samples (not contained in the simulated reference panels) in addition to the 72 trio children. We subtracted the *N*_target_=72 running times from the *N*_target_=1,072 running times to obtain the incremental cost of phasing 1,000 samples, thus adjusting for initialization costs (e.g., reading the reference data and synchronizing it with the target data), which account for a non-neglibible fraction of total computational cost when *N*_target_ is small. Finally, we divided by 1,000 to obtain per-sample costs and multiplied by 4 to scale up from the five chromosomes analyzed (≈25% of the genome) to a genome-wide analysis.

For our reference-based phasing benchmarks using GERA data, we applied an analogous procedure with the following minor differences. For each of the four sub-cohorts, we created a single simulated reference panel using all samples not in the same extended pedigree as any trio. We phased the European chip panel using Eagle1 and phased the other three panels using SHAPEIT2 (which is computationally tractable and more accurate than Eagle1 for small cohorts [13]). In each sub-cohort, we then applied each reference-based phasing method to phase all 22 autosomes of the trio parents in that sub-cohort, and we computed mean switch error rates over all trio parents. We chose to benchmark accuracy using trio parents rather than trio children due to the small numbers of trios in the non-European sub-cohorts. (For the European sub-cohort, we also computed benchmarks using trio children for comparison.) For the European sub-cohort, we computed s.e.m. over samples as before; for the other three sub-cohorts, we computed s.e.m. over 25 SNP blocks due to the small numbers of trios. When used to compare methods, these standard errors are conservative due to true variation among samples and across the genome (which causes errors to be correlated). We therefore assessed statistical significance of differences in performance between pairs of methods by performing binomial tests across samples or SNP blocks as appropriate.

**Evaluation of cohort-based phasing performance.** For our benchmarks of phasing without a reference, we created subsets of UK Biobank samples containing N=5,000, 15,000, 50,000, or 150,000 samples, each of which contained all 72 trio children and none of the 144 trio parents. We then applied each phasing method to phase chromosomes 1, 5, 10, 15, and 20 of each subset of samples, and we computed mean switch error rates and s.e.m. over the 70 European-ancestry trio children as above. We applied an analogous procedure to the GERA sub-cohorts with the same modifications as above.

## References

1. Tewhey, R., Bansal, V., Torkamani, A., Topol, E. J. & Schork, N. J. The importance of phase information for human genomics. Nature Reviews Genetics 12, 215–223 (2011).

2. Browning, S. R. & Browning, B. L. Haplotype phasing: existing methods and new developments. Nature Reviews Genetics 12, 703–714 (2011).

3. Stephens, M., Smith, N. J. & Donnelly, P. A new statistical method for haplotype reconstruction from population data. American Journal of Human Genetics 68, 978–989 (2001).

4. Halperin, E. & Eskin, E. Haplotype reconstruction from genotype data using imperfect phylogeny. Bioinformatics 20, 1842‒1849 (2004).

5. Stephens, M. & Scheet, P. Accounting for decay of linkage disequilibrium in haplotype inference and missing-data imputation. American Journal of Human Genetics 76, 449–462 (2005).

6. Scheet, P. & Stephens, M. A fast and flexible statistical model for large-scale population genotype data: applications to inferring missing genotypes and haplotypic phase. American Journal of Human Genetics 78, 629–644 (2006).

7. Browning, S. R. & Browning, B. L. Rapid and accurate haplotype phasing and missing-data inference for whole-genome association studies by use of localized haplotype clustering. American Journal of Human Genetics 81, 1084–1097 (2007).

8. Kong, A. et al. Detection of sharing by descent, long-range phasing and haplotype imputation. Nature Genetics 40, 1068–1075 (2008).

9. Browning, B. L. & Browning, S. R. A unified approach to genotype imputation and haplotype-phase inference for large data sets of trios and unrelated individuals. American Journal of Human Genetics 84, 210–223 (2009).

10. Delaneau, O., Marchini, J. & Zagury, J.-F. A linear complexity phasing method for thousands of genomes. Nature Methods 9, 179–181 (2012).

11. Williams, A. L., Patterson, N., Glessner, J., Hakonarson, H. & Reich, D. Phasing of many thousands of genotyped samples. American Journal of Human Genetics 91, 238–251 (2012).

12. Delaneau, O., Zagury, J.-F. & Marchini, J. Improved whole-chromosome phasing for disease and population genetic studies. Nature Methods 10, 5–6 (2013).

13. Loh, P.-R., Palamara, P. F. & Price, A. L. Fast and accurate long-range phasing in a uk biobank cohort. Nature Genetics (2016).

14. O’Connell, J. et al. Haplotype estimation for biobank-scale data sets. Nature Genetics (2016).

15. Snyder, M. W., Adey, A., Kitzman, J. O. & Shendure, J. Haplotype-resolved genome sequencing: experimental methods and applications. Nature Reviews Genetics 16, 344–358 (2015).

16. van de Geijn, B., McVicker, G., Gilad, Y. & Pritchard, J. K. WASP: allele-specific software for robust molecular quantitative trait locus discovery. Nature Methods 12, 1061–1063 (2015).

17. Kumasaka, N., Knights, A. J. & Gaffney, D. J. Fine-mapping cellular QTLs with RASQUAL and ATAC-seq. Nature Genetics (2015).

18. McCarthy, S. *et al.* A reference panel of 64,976 haplotypes for genotype imputation. bioRxiv 035170 (2016).

19. Durbin, R. Efficient haplotype matching and storage using the positional Burrows-Wheeler transform (PBWT). Bioinformatics 30, 1266–1272 (2014).

20. Li, N. & Stephens, M. Modeling linkage disequilibrium and identifying recombination hotspots using single-nucleotide polymorphism data. Genetics 165, 2213–2233 (2003).

21. Sudlow, C. et al. UK Biobank: an open access resource for identifying the causes of a wide range of complex diseases of middle and old age. PLOS Medicine 12, 1–10 (2015).

22. Kvale, M. N. et al. Genotyping informatics and quality control for 100,000 Subjects in the Genetic Epidemiology Research on Adult Health and Aging (GERA) Cohort. Genetics 200, 1051–1060 (2015).

23. Banda, Y. et al. Characterizing race/ethnicity and genetic ancestry for 100,000 subjects in the Genetic Epidemiology Research on Adult Health and Aging (GERA) cohort. Genetics 200, 1285–1295 (2015).

24. Browning, B. L. & Browning, S. R. Genotype imputation with millions of reference samples. The American Journal of Human Genetics 98, 116–126 (2016).

25. 1000 Genomes Project Consortium et al. A global reference for human genetic variation. Nature 526, 68–74 (2015).

26. Howie, B., Fuchsberger, C., Stephens, M., Marchini, J. & Abecasis, G. R. Fast and accurate genotype imputation in genome-wide association studies through pre-phasing. Nature Genetics 44, 955–959 (2012).

27. He, D., Han, B. & Eskin, E. Hap-seq: an optimal algorithm for haplotype phasing with imputation using sequencing data. Journal of Computational Biology 20, 80–92 (2013).

28. Delaneau, O., Howie, B., Cox, A. J., Zagury, J.-F. & Marchini, J. Haplotype estimation using sequencing reads. American Journal of Human Genetics 93, 687–696 (2013).

29. Sharp, K., Kretzschmar, W., Delaneau, O. & Marchini, J. Phasing for medical sequencing using rare variants and large haplotype reference panels. Bioinformatics (2016).

30. Chang, C. C. et al. Second-generation PLINK: rising to the challenge of larger and richer datasets. GigaScience 4, 1–16 (2015).

31. Browning, S. R. Multilocus association mapping using variable-length Markov chains. American Journal of Human Genetics 78, 903–913 (2006).

32. Browning, B. L. & Browning, S. R. Efficient multilocus association testing for whole genome association studies using localized haplotype clustering. Genetic Epidemiology 31, 365–375 (2007).

33. McVean, G. A. & Cardin, N. J. Approximating the coalescent with recombination. Philosophical Transactions of the Royal Society of London B: Biological Sciences 360, 1387–1393 (2005).

34. Palamara, P. F., Lencz, T., Darvasi, A. & Pe’er, I. Length distributions of identity by descent reveal fine-scale demographic history. American Journal of Human Genetics 91, 809–822 (2012).

35. Harris, K. & Nielsen, R. Inferring demographic history from a spectrum of shared haplotype lengths. PLOS Genetics 9, e1003521 (2013).

36. Drmanac, R. et al. Human genome sequencing using unchained base reads on self-assembling DNA nanoarrays. Science 327, 78–81 (2010).

37. Huang, J. et al. Improved imputation of low-frequency and rare variants using the UK10K haplotype reference panel. Nature Communications 6 (2015).

